# NEUROTRANSMITTER TRANSPORTER/RECEPTOR CO-EXPRESSION SHARES ORGANIZATIONAL TRAITS WITH BRAIN STRUCTURE AND FUNCTION

**DOI:** 10.1101/2022.08.26.505274

**Authors:** Benjamin Hänisch, Justine Y. Hansen, Boris C. Bernhardt, Simon B. Eickhoff, Juergen Dukart, Bratislav Misic, Sofie L. Valk

## Abstract

The relationship between brain areas based on neurotransmitter receptor and transporter molecule expression patterns may provide a link between brain structure and its function. Here, we studied the organization of the receptome, a measure of regional neurotransmitter receptor/transporter molecule (NTRM) similarity, derived from in vivo PET imaging studies of 19 different receptors and transporters. Nonlinear dimensionality reduction revealed three main spatial gradients of receptor similarity in the cortex. The first gradient differentiated the somato-motor network from the remaining cortex. The second gradient spanned between temporo-occipital and frontal anchors, differentiating visual and limbic networks from attention and control networks, and the third receptome gradient was anchored between the occipital and temporal cortices. In subcortical structures, the receptome delineated a striato-thalamic axis, separating functional communities. Moreover, we observed similar organizational principles underlying receptome differentiation in cortex and subcortex, indicating a link between subcortical and cortical NTRM patterning. Overall, we found that the cortical receptome shared key organizational traits with brain structure and function. Node-level correspondence of receptor similarity to functional, microstructural, and diffusion MRI-based measures decreased along a primary-to-transmodal gradient. Compared to primary and paralimbic regions, we observed higher receptomic diversification in unimodal and heteromodal regions, possibly supporting functional flexibility. In sum, we show how receptor similarity may form an additional organizational layer of human brain architecture, bridging brain structure and function.

## Introduction

Uncovering how the anatomy of the brain supports its function is a long-standing goal of neuroscientific research^1^. Neuroanatomical mapping of cyto- and myeloarchitecture^2–4^, in combination with lesion studies^5–7^ have established that there is substantial variability in cellular composition of brain areas and a relationship between a brain area’s structure and its function. Neurotransmitter receptor distribution is an important additional layer of brain organization. Neurotransmitters and their respective receptors convey neurotransmission, an essential aspect of neural communication. As such the biologically versatile nature of neurotransmitter-receptor interactions is a pivotal component of a neuronal system^8^. To understand the role of neurotransmission in brain functionality, deciphering the neurotransmitter receptor distribution landscape is thus essential.

Autoradiography studies have shown that neurotransmitter receptors are heterogeneously expressed throughout the cortex. Generally, receptor distributions vary in a horizontal laminar fashion similar to cyto- and myeloarchitectural cortical layers^9,10^, but are also closely related to vertical cyto- and myeloarchitectural cortical composition. Receptor distributions recapitulate histology-defined cortical areas, but can also group different cortical areas into neurochemical families as well as further subdivide cytoarchitecturally homogeneous regions^10,11^. Changes in localized brain function are reflected by changes in receptor distributions, demonstrated for example in the changes of multiple receptor densities at the border between primary (V1) and secondary (V2) visual cortex^12,13^. Furthermore, a similarity in receptor architecture can be observed between brain areas sharing similar functionality. This suggests receptor “fingerprints”, the density profiles of multiple neurotransmitter receptor types in a specific brain area, as key features of functional specialization^10,13–15^. Receptor fingerprints delineate sensory from association cortices^16^, and provide a common molecular basis of areas involved in language comprehension^17^. Studies of receptors via autoradiography have shown links between receptor fingerprints and brain functionality in localized brain areas. However, a coherent whole-brain perspective on receptor profile similarities and their relationship to brain function is missing. Importantly, a whole-brain perspective should include subcortical structures. Although the subcortical distributions of some receptors have been systematically investigated^18–20^, there are no systematic studies of receptor co-distributions or similarity in the subcortex. Characterizing subcortical receptor similarity profiles could be an important step towards the understanding of this so far understudied part of the human brain^21^.

A whole-brain perspective on receptor similarity could provide a promising avenue towards understanding brain functionality, as the interaction of different neurotransmitter systems is a major component of any functional process. This is reflected by the reorganization of functional brain networks after pharmacological perturbation of neurotransmitter systems^22–24^. This has led to the hypothesis that neuromodulation through neurotransmission might play a major role in enabling the static nature of brain structure to yield flexible functionality^25,26^. Functional co-activations are only partially explained by the physical connectedness of brain areas^1,27^, hinting at considerable relevance of other aspects of brain organization to understand structure-function relationships. Incorporating neurotransmitter co-distribution as an additional layer of brain architecture thus holds promise of further insights into the structural basis of brain function.

Moreover, understanding how the organization of receptor distributions relates to brain functionality holds clinical relevance. An extensive body of research links alterations in neurotransmitter receptor and transporter expression patterns to psychiatric diseases^28–31^. Most psychotropic drugs manipulate the brain’s neurotransmission landscape and are, although their mechanisms of action are often incompletely understood, effective and reliable pillars in the treatment of psychiatric diseases^32–35^. However, there is no governing rule that ties neurotransmission to distinct aspects of cerebral dysfunction. Historical perspectives linking single neurotransmitters to mental illness^36,37^ have gradually shifted towards the understanding that clinical phenotypes can be associated with alterations in multiple neurotransmitter systems^38–40^. The study of receptor co-expression could thus provide novel avenues towards understanding the neurobiology of psychiatric diseases^41–44^. This leaves us with two main questions; 1) How is similarity in receptor fingerprints organized in the cortex and subcortical structures? and 2) How does similarity in receptor fingerprints relate to structural features of brain organization and different aspects of brain function?

The whole-brain spatial distribution of neurotransmitter transporter and receptor molecule (NTRM) densities in humans can be measured using Positron-Emission Tomography (PET). Indeed, recent work used PET-derived NTRM density measurements to elucidate the role of different receptor profiles in mediating structure-function relationships and functional processing, providing novel insights into the role of neurochemical profiles in brain architecture^45^. However, the interrelationship of brain areas based on their neurochemical similarity profiles, and which organizational principles support neurotransmitters to convey brain functionality, remain underexplored^17^. Here, we capture the interrelationship of brain areas based on their receptor fingerprints. We calculate the covariance network of 19 regional NTRM densities across 1200 subjects, which we term the ‘receptome’, from a dataset previously compiled by Hansen et al^45^. We first study how receptor similarity across the cortex and subcortical structures is spatially organized. For this, we employ an unsupervised dimensionality reduction technique to generate principal gradients, which are low-dimensional representations of the organizational axes in the cortical and subcortical receptome. Using these gradients, we identify NTRM distributions that drive regional receptor (dis)similarity. Several follow-up analyses shed light upon the relationship to organizational axes in structural connectivity (SC)^46^, Microstructural Profile Covariance (MPC)^47^ and resting-state functional connectivity (rsFC)^48^, as well as to term-based functional brain activation^49^ and radiological markers of disease^50^. Finally, we performed various analyses to evaluate robustness of our observations.

## Results

### Organization of the cortical receptome (Figure 1)

To assess cortical receptor similarity, we leveraged a large publicly available dataset of PET-derived NTRM densities, containing 19 different NTRM from a total of over 1200 subjects^45^. After parcellation^51^, we calculated a Spearman rank correlation matrix of parcel-level NTRM densities, the receptome (**Fig. 1A**). The receptome represents node-level interregional similarities in receptor fingerprints. We next employed principal gradient decomposition to delineate the main organizational axes of cortical receptor similarity (**Fig. 1A, Methods**). We identified three principal gradients, explaining 15%, 14% and 13% of variance, respectively (**Fig. 1A**). The first receptome gradient (RC G1) described an axis stretching between sensory-motor regions and inferior temporal and occipital lobe. The second receptome gradient (RC G2) spanned between a temporo-occipital and a frontal anchor. Finally, the third receptome gradient (RC G3) was differentiated between the occipital cortex and the temporal lobe (**Fig. 1B**).

**Fig. 1.**
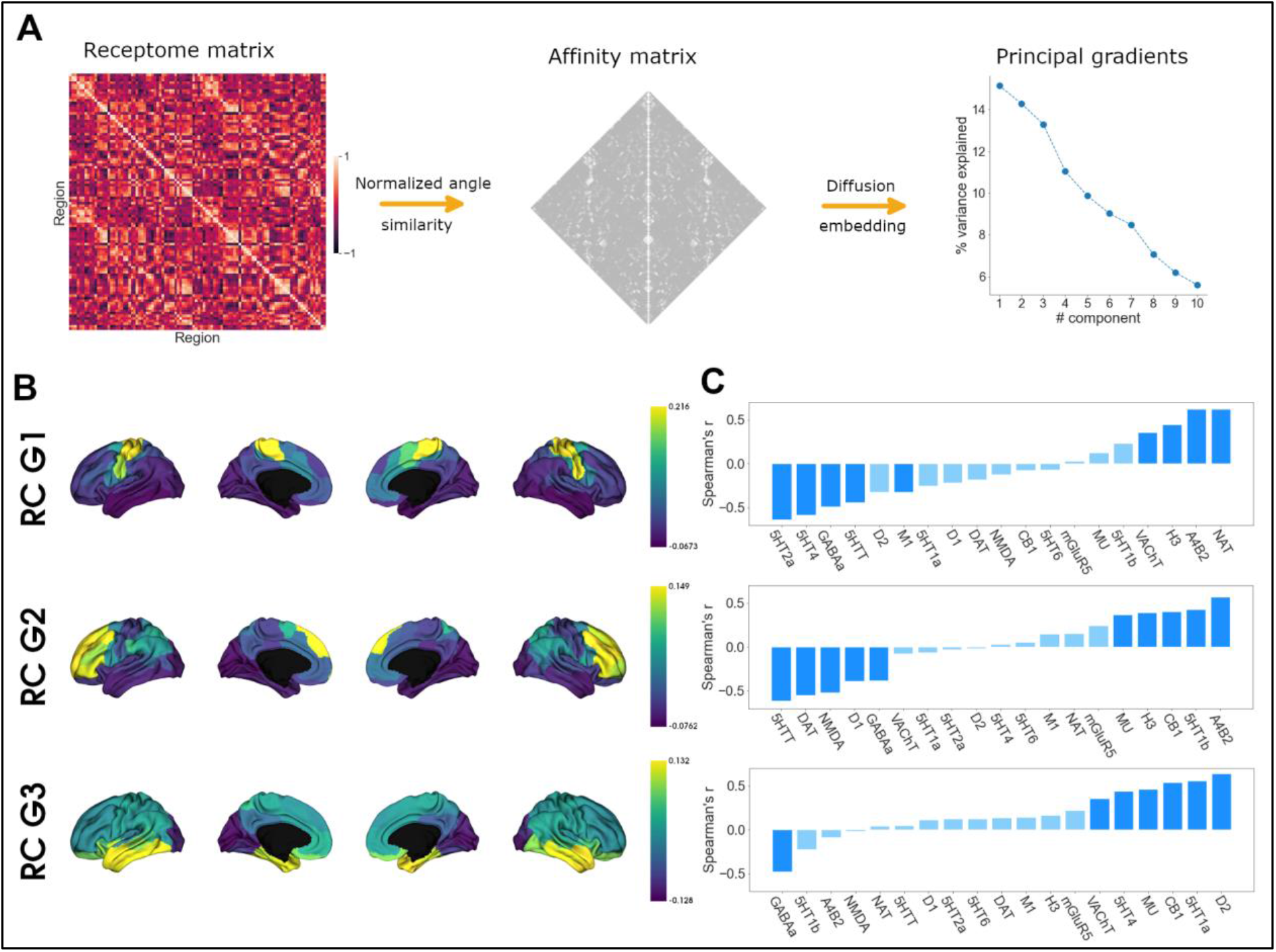
Organization of the cortical receptome. **A)** Analytic workflow of receptome principal gradient decomposition. Spearman rank correlation captures node-level similarity in chemoarchitectural composition, generating the receptome matrix. Next, to determine similarity between all rows of the receptome matrix, we used a normalized angle similarity kernel to generate an affinity matrix. Finally, we employ diffusion embedding, a nonlinear dimensionality reduction technique, to derive principal gradients of receptomic organization. **B)** Receptome gradients projected on the cortical surface. Top: First receptome gradient; Middle: Second receptome gradient; Bottom: Third receptome gradient. **C)** Spearman rank correlations of principal receptome gradients with individual NTRM densities. Top: First receptome gradient; Middle: Second receptome gradient; Bottom: Third receptome gradient.

To determine which NTRM distributions drive the main axes of cortical receptor similarity, we performed Spearman rank correlations between a parcel’s associated gradient value and its receptor fingerprint, meaning the density of each receptor/transporter molecule in it (**Fig. 1C**). Note that the gradient value of a parcel is a measure of where on the gradient axis the parcel is located, from which similarity to parcels with similar values, and dissimilarity to parcels with dissimilar values, is inferred. Thus, a receptor that is most strongly expressed in parcels with negative values, while having weak expression in parcels with positive values, will be negatively correlated to the gradient. RC G1 was primarily driven by the anticorrelation between distributions of 5-HTT, 5-HT4, 5-HT2a and GABAa with the distributions of VAChT, H3, NAT and Α4Β2. RC G2 separated 5-HTT, DAT, NMDA, D1 and GABA distributions from α4β2, 5-HT1b, CB1, H3 and MU. RC G3 showed significant negative correlations to GABAa distributions and significant positive correlations to D1, 5-HT1a, CB1, MU, 5-HT4 and VAChT.

### Organization of the subcortical receptome (Figure 2)

After we delineated main axes of variation in cortical receptor similarity, we investigated how NTRM co-distribution is organized across subcortical structures. We selected the caudate nucleus, putamen, nucleus accumbens, pallidal globe, amygdala and thalamus as regions of interest, since they are well characterized as well as of reasonable size to be investigated by PET imaging. Due to different tracer uptake dynamics in the cortex and subcortical structures, we analyzed the subcortex separately from the cortico-cortical PET covariance.

**Fig 2.**
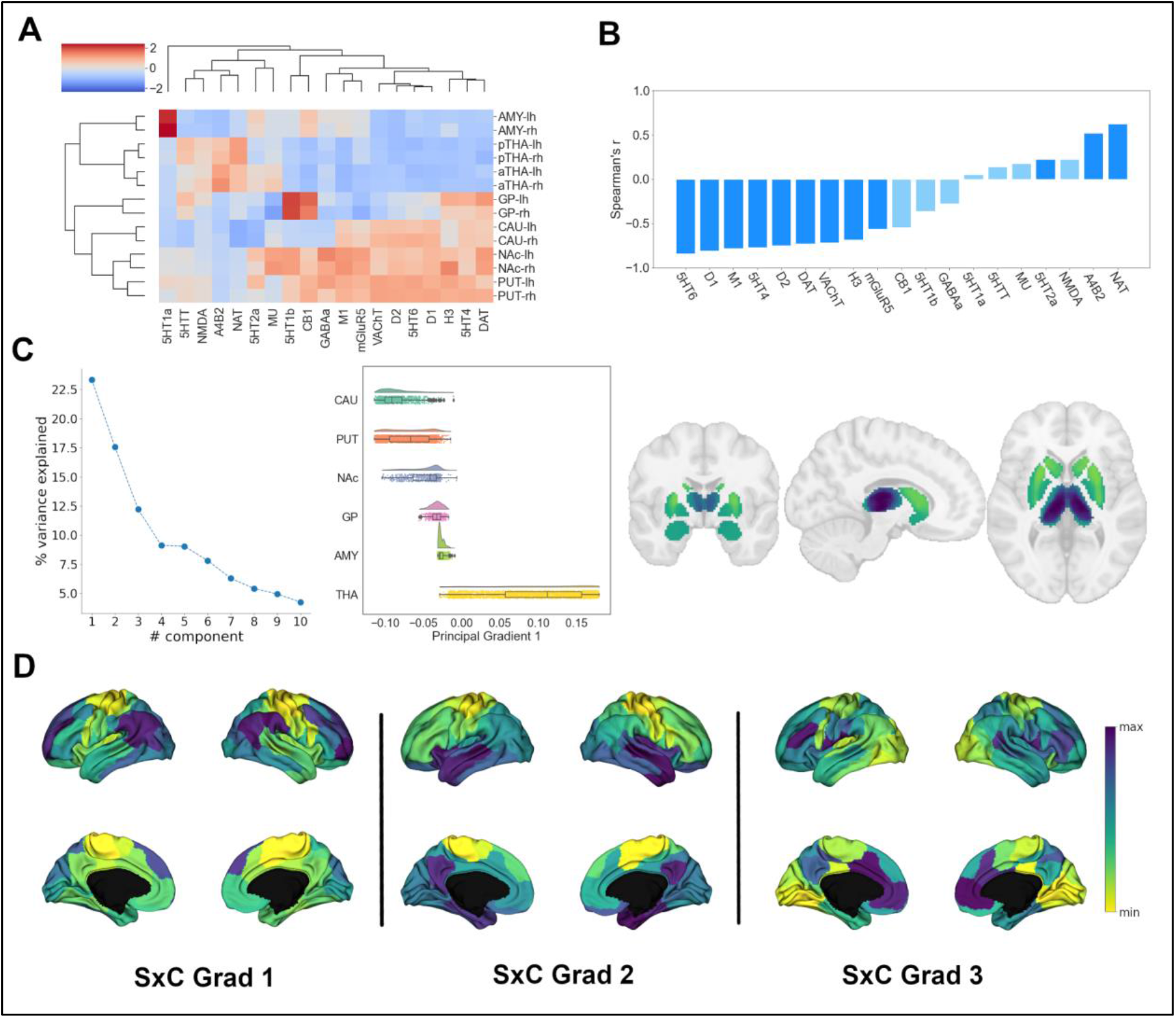
Organization of the subcortical receptome. **A)** Hierarchical agglomerative clustering of NTRM densities in subcortical structures. aTHA: anterior thalamus; pTHA: posterior thalamus. **B)** Spearman rank correlations of the first subcortical principal receptome gradient with individual NTRM densities **C)** Principal receptome gradient decomposition of the subcortical receptome. *Left:* Percentage of variance explained by components following principal gradients decomposition. *Middle:* Distribution of values of the first principal gradient of the subcortical receptome across subcortical structures. CAU: caudate nucleus; PUT: putamen; NAc: accumbens nucleus; GP: pallidal globe; AMY: amygdala; THA: thalamus. *Right:* Subcortical projection of the first principal gradients of the subcortical receptome **D)** Principal gradients of the subcortico-cortical receptome projected to the cortical surface.

First, we were interested in how receptor fingerprints differentiate subcortical structures, and which NTRM co-distributions drive this separation. We thus performed agglomerative hierarchical clustering on the z-scored mean NTRM density of all subcortical structures per hemisphere (**Fig. 2A**). We found that subcortical NTRM expression was largely symmetrical between hemispheres, as indicated by the immediate clustering of structures with their counterpart from the other hemisphere. The main hierarchical branch separated putamen, accumbens nucleus, caudate nucleus and pallidum from amygdala and thalamus. Putamen, accumbens nucleus and caudate nucleus make up the striatum, and the three structures displayed high similarity in receptor fingerprints. We found thalamus and striatum to have considerable differences in NTRM co-expression patterns. α4β2, NAT, 5-HTT and NMDA showed strong co-expression in thalamus but not in striatum, while D1, D2, DAT, 5-HT4, 5-HT6, M1 and VAChT were strongly co-expressed in striatum, but not in thalamus.

After investigating individual receptor fingerprints, we analyzed receptor similarity in subcortical structures. We performed voxel-wise Spearman rank correlations of subcortical NTRM densities to construct a subcortical receptome. To discern how subcortical substructures can be reconstructed based on receptor similarity, we employed the Leiden community detection method^52^, a greedy optimization algorithm that opts to minimize variance within and maximize variance between communities. Subcortical receptome clustering exhibited high stability across the resolution parameter sample space (**Fig. S1A**). Receptomic clustering discerned three dominant communities, the first majorly capturing the striatal structures (putamen, caudate, NAc) and the pallidal globe, the second majorly capturing the thalamus, and the third majorly capturing the amygdala (**Fig. S1A)**. We then used diffusion embedding to derive low-dimensional gradient embeddings of the subcortical receptome to discern its main organizational axes. The first subcortical receptome gradient (sRC G1) explained 23% of variance. It extended between the striatum (caudate nucleus, putamen) across the pallidal globe to the thalamus, with the amygdala being in the middle (**Fig. 2C**). Note that proximity of structures was not a major determinant of sRC G1 values, demonstrated by voxels of the caudate nucleus and thalamus that are located closely to each other but showed diverging sRC G1 values. The second gradient, explaining 17.5% of variance, and third gradient, explaining 12% of variance, described a ventral-dorsal and medial-lateral trajectory, respectively (**Fig. S1D, S1E**). To determine the influence of different subcortical NTRM distributions on subcortical receptome gradients, we performed voxel-wise Spearman rank correlations between gradient values and receptor fingerprints. The first subcortical receptome gradient showed significant positive correlations to NAT, α4β2 and 5-HT2a densities, and significant negative correlations to 5-HT6, D1, M1, 5-HT4, D2, DAT, VAChT, H3 and mGluR5 distributions (**Fig. 2D**).

Last, we were interested in the relationship between the subcortical and cortical receptomes. We constructed a subcortico-cortical NTRM covariance matrix and subsequently applied diffusion embedding to delineate the principal gradients of subcortico-cortical receptor similarity (**Fig. 2F**). To quantify the overlap between cortical and subcortico-cortical main organizational principles, we performed Spearman rank correlations between cortical and subcortico-cortical receptome gradients. The first and second cortical gradients correlated significantly with all subcortico-cortical receptome gradients, while the third cortical gradient only correlated significantly to the third subcortico-cortical gradient (**Fig. S1C**).

### Relationship of the cortical receptome to brain function and disease (Figure 3)

After characterizing the cortical and subcortical receptomes, we next sought to investigate the relationship of receptor similarity to hallmarks of brain function and dysfunction. To dissect the relationship between cortical receptor similarity and brain function, we used term-based meta-analytical maps of functional brain activation. This approach associates a functional term with localized brain activity (e.g. ‘primary somatomotor’ is associated with activation in the precentral gyrus). Using the neurosynth database^49^, we calculated Spearman rank correlations between normalized activation maps and receptor similarity gradients (**Fig. 3B**). We selected activation maps based on a curated list of terms of interest to cover multiple dimensions of term-related functional activation. Briefly, we selected for a range of terms associated with unimodal (e.g. sensory-motor) to transmodal (e.g. information integration, emotion processing, social cognition) functionality, as well as for terms of neurological and psychiatric diseases (e.g. dementia). The full list can be found in **Supplement L1**. Negative correlations imply a relationship between a term-based functional activation mainly located in parcels with negative gradient values. RC G1 showed strong positive correlations with meta-analytical terms related to somato-motor function, followed by terms related to cognitive control and abstract terms (e.g. ‘illusion’, ‘insight’). Its strongest significant negative correlations were to terms related to visual function, memory and Theory of Mind. Functional decoding of RC G2 revealed a processing hierarchy from primary visual terms to terms of complex cognitive functions best subsumed under control, cognitive constructs, decision making, memory and (social) cognition, but also sensory-motor terms. Functional decoding of RC G3 revealed a differentiation of terms related to visual processing from terms related to auditory processing, social cognition and memory.

**Fig 3.**
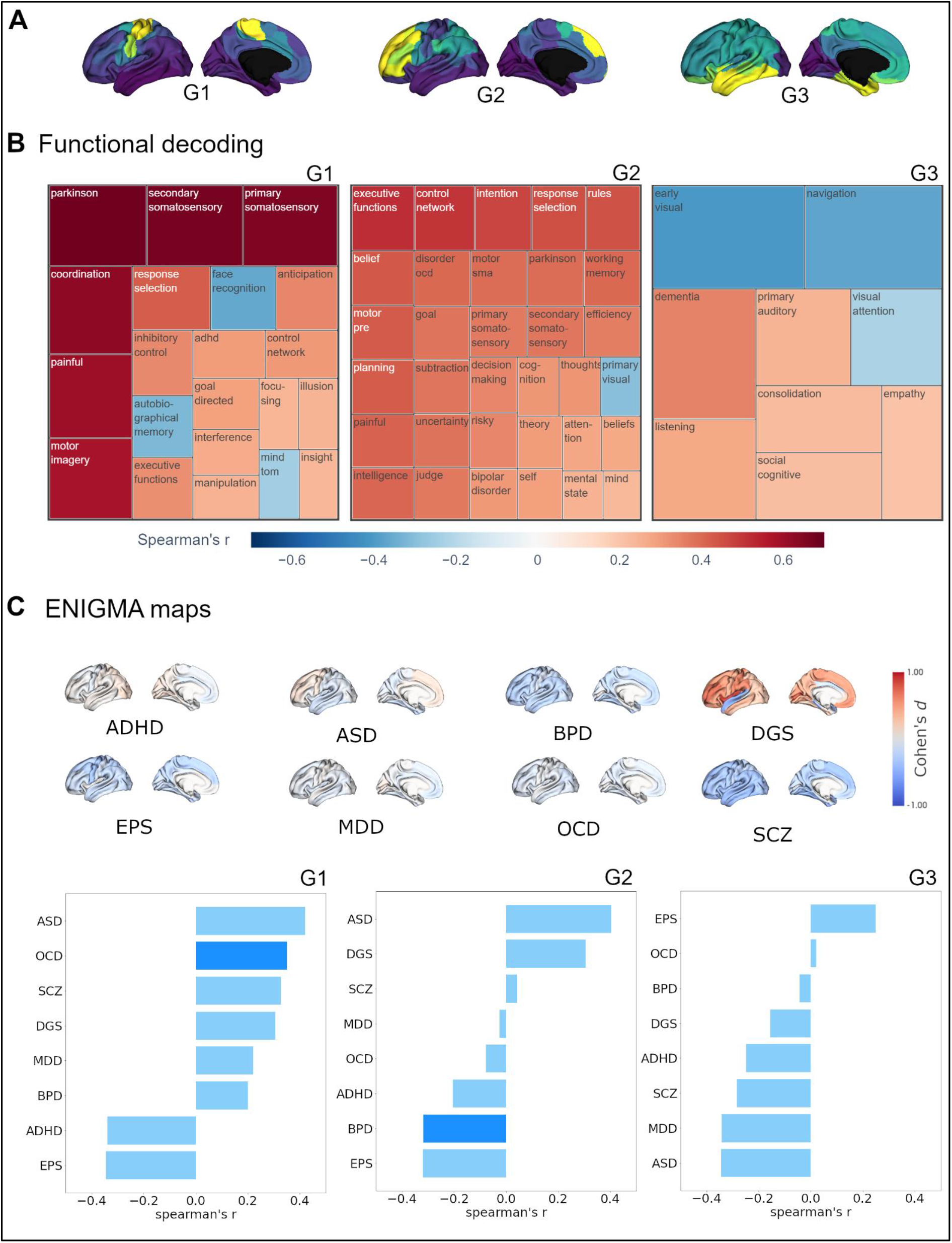
Cortical receptome gradients in task-based functional activation and disorder. **A)** Cortical receptome gradients projected to the cortical surface. **B**) Functional decoding of cortical receptome gradients. Treemaps display positive and negative correlations of receptome gradients and term-based functional activation patterns. Rectangle sizes encode absolute correlation strength. Note that the coloring in all treemaps encodes the same correlation values, while rectangle sizes are better suited to compare the within-gradient relevance of terms. Left: RC G1; Middle: RC G2; Right: RC G3. **C)** Disease decoding of cortical receptome gradients. Surface plots: Effect size (Cohen’s d) of cortical thickness alterations in central nervous system disorders in patients vs controls. ASD: Autism Spectrum Disorder; ADHD: Attention Deficit Hyperactivity Disorder; BPD: Bipolar Disorder; DGS: DiGeorge-Syndrome (22q11.2 deletion syndrome); EPS: Epilepsy; MDD: Major Depressive Disorder; OCD: Obsessive-Compulsive Disorder; SCZ: Schizophrenia. Bar plots: Spearman rank correlations of receptome gradients and cortical thickness alterations. Left: RC G1; Middle: RC G2; Right: RC G3.

To assess the role of the cortical receptome in dysfunctional brain states, we chose to compare receptome gradients to structural brain abnormalities in psychiatric and neurological disorders. We leveraged disease-related variations in cortical thickness, a radiological marker of structural brain abnormalities, derived via a standardized multi-site effort^50^. Cortical thickness was quantified by Cohen’s d case-vs-control effect size and accessed through the ENIGMA toolbox^53^. We chose autism spectrum disorder (ASD)^54^, attention deficit hyperactivity disorder (ADHD)^55^, bipolar disorder (BPD)^56^, DiGeorge-syndrome (22q11.2 deletion syndrome) (DGS)^57^, epilepsy (EPS)^58^, major depressive disorder (MDD)^59^, obsessive compulsive disorder (OCD)^60^ and schizophrenia (SCZ)^61^ to cover a broad spectrum of diseases (**Fig. 3C**). Receptome gradients captured disease-specific cortical thickness alteration patterns. RC G1 showed significant positive correlations to the cortical thickness profile of obsessive-compulsive disorder, while RC G2 had significant negative correlations to cortical thickness alterations in bipolar disorder. Both OCD and BPD are primarily associated with cortical thinning, thus, cortical thickness in OCD is significantly reduced where RC G1 values are positive, and significant reductions in BPD are where RC G2 values are negative. RC G3 does not show significant associations with cortical disease profiles. (**Fig. 3C**).

### Interrelationship between the cortical receptome and structural, functional and cytoarchitectural organization (Figure 4)

Finally, we investigated the relationship of cortical receptor similarity to other measures of whole-brain organization. As autoradiography studies connect receptor distributions to cytoarchitectural characteristics^10^, we studied the relationship of cortical receptomic organization to Microstructural Profile Covariance (MPC), an MRI-derived proxy measure of regional myelin content similarity that also reflects cytoarchitectural variations^62^, and a gradient of cytoarchitectural variation derived from the BigBrain project^47,63^ (BB G1). Additionally, to further investigate structural and functional relevance, we explored the relationships of cortical receptor similarity to diffusion MRI tractography-derived structural connectivity (SC), and functional MRI-derived resting-state functional connectivity (FC).

**Fig. 4.**
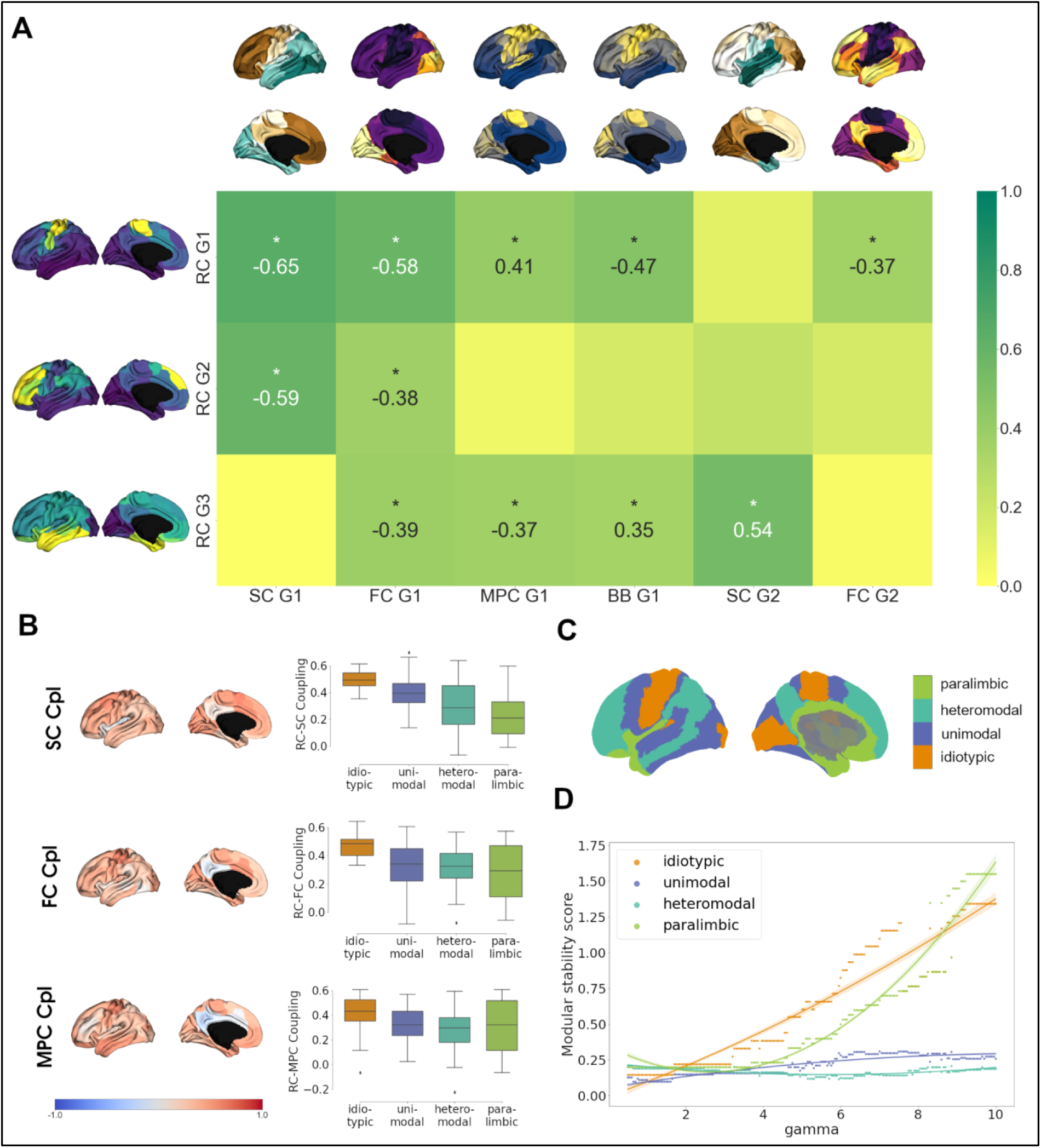
Cortical NTRM covariance. **A)** Heatmap of correlation strengths between cortical receptome gradients and FC, SC and MPC gradients and the BigBrain gradient (BB G1). The color scheme is scaled to absolute values. Gradients are displayed on the cortical surface next to their respective rows and columns **B)** Coupling of the cortical receptome to SC, FC and MPC. *Left*: Projection of coupling strengths on the cortical surface. *Right*: Coupling strengths by cytoarchitectural classes. **C)** Mesulam cytoarchitectural classes projected to the cortical surface **D)** Modular stability of receptome similarity clustering in Mesulam cytoarchitectural classes, reflecting the heterogeneity of receptomic profile.

We first chose to compare receptome gradients to gradients of SC, FC and MPC, reflecting principal organizational axes within these modalities. Due to the amount of variance the respective gradients explain, we focused on the first two principal gradients of SC and FC, and the first principal gradient of MPC **(Fig. S2A)**. Additionally, we analyzed the relationship of receptome gradients to networks of functional connectivity (**Fig. S2C**)^64^. RC G1 showed strongest overlaps to SC G1 and FC G1, as these gradients shared either anterior-posterior or visual-to-somatomotor trajectories **(Fig. 4A)**. Additional significant, but weaker correlations for RC G1 were to BB G1 and MPC G1, which represent the main axes of cortical cytoarchitectural similarity^47^, and FC G2, which separates unimodal from association cortices^65^. Functional network decoding revealed that RC G1 separates visuo-limbic from somatomotor cortices. Similar to the first receptome gradient, RC G2 correlated significantly to SC G1 and FC G1, while separating visuo-limbic from control networks. RC G3 showed the strongest correlations to SC G2, which separated occipital and temporal cortex. Further significant correlations were to FC G1, MPC G1 and BB G1. While there was a close relationship between visual and limbic networks in the first two receptome gradients, functional network decoding placed these networks on opposite ends on RC G3.

After finding overlaps in main organizational axes, we next investigated node-level similarities between the receptome and FC, SC and MPC. We performed row-wise correlations of the receptome matrix to each other aforementioned matrix, respectively. Taking SC as an example, this yielded one value per parcel expressing the similarity between a parcel’s receptomic relationship to every other parcel and its structural connectivity relationship to every other parcel (**Fig. 4B**). The resulting correlation coefficients expressed the strength of coupling between two measures. Generally, coupling strength of the receptome to the aforementioned measures was found to decrease along a sensory-fugal gradient of laminar differentiation, an influential theoretical framework that attributes cognitive processing complexity to cortical areas using cytoarchitectural classes^66^. Average coupling strength across cytoarchitectural classes was significantly different across all metrics. RC-SC decoupling along the sensory-fugal gradient (Kruskal-Wallis’ *h*=24.43, *p* < 0.001) was driven by significantly stronger coupling in idiotypic relative to heteromodal and paralimbic cortices (post-hoc Dunn’s test with Bonferroni correction *p* < 0.001). RC-FC coupling strengths in idiotypic cortices were significantly increased relative to unimodal, heteromodal and paralimbic cortices (*h*=16.68, *p* < 0.001; Dunn’s test *p* < 0.02). Last, RC-MPC decoupling across cytoarchitectural classes (*h*=9.16, *p* < 0.05) was primarily reflected by decreased coupling in heteromodal versus idiotypic regions (Dunn’s test *p* < 0.02).

Our previous decoding results hint at a relationship between cortical hierarchy and receptomic characteristics. We thus aimed to analyze cortical receptomic heterogeneity in the context of cytoarchitectural classes^66^. To this end, we leveraged the Leiden community detection algorithm to discover cortical communities of receptor similarity. We observed that new communities primarily formed in the frontal cortex when sampling the parameter space, indicating more unique receptor fingerprints. To capture how stably receptomic communities recapitulate cytoarchitectural classes when increasing the number of receptomic communities detected, we developed the modular stability score (see **Methods**). A cytoarchitectural class that is largely covered by a single receptomic community and does not increasingly fracture with an increase in the overall number of communities has a high modular stability score. Overall, paralimbic cortices exhibited modular stability similar to idiotypic cortices, while heteromodal and unimodal regions were less stable (**Fig. 2D**). This suggests that idiotypic and paralimbic cortices show a more homogeneous receptomic profile, while heteromodal and unimodal cortices have a more diverse chemoarchitectural landscape. We made similar observations studying the relationship of receptomic communities to networks of resting-state functional connectivity^64^, where visual, limbic and sensory-motor networks exhibited higher modular stability than the ventral and dorsal attention, control and default mode networks (**Fig. S2C**). In further investigation through agglomerative hierarchical clustering of mean NTRM densities in functional networks, we found that dorsal and ventral attention, somato-motor and default networks were first combined into one cluster, before visual and finally limbic networks were merged to that cluster (**Fig. S2D**). This indicates more distinct chemoarchitectural profiles in limbic and visual networks with little overlap amongst each other.

### Robustness analysis

Owing to the low spatial resolution of PET NTRM imaging, we chose to present our main findings in the coarse resolution of 100 Schaefer parcels. To assess validity, we replicated our analyses in Schaefer parcellations 200-400^51^. Selecting a finer granularity than 400 parcels was not reasonable due to the limited resolution of PET images^67^. Surface projections of receptome gradients showed good replicability across parcellations (**Fig. S3A-D**), although an increase in parcellation granularity shifted one extreme in RC G1 and RC G2 towards the temporal poles. Notably, for granularities of 200 and 400 parcels, there is a component ranking switch meaning that the pattern captured by RC G1 in the main results is captured by RC G2 in the replication, and vice-versa. As gradients of rsFC, SC and MPC also change as a function of parcellation granularity, we repeated the correlation analyses across different parcellations. The shift towards the temporal pole in RC G1 and G2 lead to a clearer separation between one receptome gradient that strongly correlated to SC G1, and another one that significantly correlated to FC G2 in parcellation granularities 200 and 300 (**Tables S2A-D**). Additionally, to ensure robustness of hierarchical clustering results of subcortical and cortical NTRM densities, we replicated the analysis using different linkage methods (**Fig. S4, Fig. S5**).

## Discussion

In the present work, we set out to investigate the organizational principles of neurotransmitter transporter and receptor similarity in the human cerebral cortex and subcortex. Additionally, we aimed to dissect the functional relevance of receptor similarity and its relationship to other measures of brain organization. Thus, we studied the connection of cortical receptor similarity to markers of brain function and disease, and explored its relationship to the structural, functional and cytoarchitectural organization of the cortex. In sum, we introduce and thoroughly characterize receptor similarity as an additional layer of macro-scale brain architecture. Leveraging this architectural layer, we present novel insights into structure-function relationships in the human brain.

A cornerstone technique of our study was the use of a nonlinear dimensionality reduction technique to derive principal gradients of the receptome, a matrix of regional receptor similarity. The first receptome gradient, RC G1, describes an axis stretching between sensory-motor regions and inferior temporal and occipital lobe, and differentiates the density profiles of 5-HT2a, 5-HT4, GABAa, 5-HTT and M1 from VAChT, H3, α4β2 and NAT. On the organizational level, the first axis of cortical receptor similarity combines key features of structural and functional organization. RC G1 established similar relationships between cortices as the organization of physical connections, captured by SC G1, which is likely driven by the distance-dependent nature of cortical wiring^68^. It also captures meaningful variation in cytoarchitecture, exemplified by correlations MPC G1 and BB G1, and functional organization, signified by correlations to FC G1 and FC G2, although these correlations are not consistent across parcellation granularities. The anchors of RC G1 on the one end are involved in somatomotor and control functions, and visual, memory and socio-cognitive functions (Theory of Mind) on the other end, as revealed by term-based activation decoding. Finally, RC G1 correlates significantly with cortical thickness alterations patterns associated with obsessive-compulsive disorder. Taken together, the first receptome gradient can be summarized to primarily capture the differentiation of the somato-motor network from the remaining cortex, with the most pronounced differences outlined against visual and limbic cortices. This dissimilarity is likely linked on one side by the differences in expression patterns of 5-HTT, 5-HT4, 5-HT2a, GABAa and M1, which are predominantly expressed in the temporal and occipital cortices. On the other side, NAT, α4β2, H3 and VAChT are predominantly expressed pericentrally and in the frontal cortex. RC G1 also connects these receptor co-expression profiles to morphological changes in obsessive-compulsive disorder, where the relationship it outlines to serotonin signaling is particularly interesting. Selective Serotonin Reuptake Inhibitors (SSRIs) target 5-HTT and are the preferred pharmacological intervention in the treatment of OCD^34,69^. Genetically, 5-HT2a and 5-HTT variants have been identified as risk factors for the development of obsessive-compulsive disorder^70^, and OCD patients showed in peripheral 5-HTT and 5-HT2a functionality^71^. Furthermore, there is emerging evidence that GABA signaling abnormalities are related to the development of obsessive-compulsive disorder^44^, although conclusive evidence is lacking.

The second gradient of receptor similarity, RC G2, spans between temporo-occipital and frontal anchors, separating visual and limbic networks from attention and control networks. This gradient separates 5-HTT, DAT, NMDA, D1 and GABAa from MU, H3, CB1, 5-HT1b and α4β2. It correlates significantly to the first gradients of structural and functional connectivity. Term-based functional activation decoding reveals that RC G2 spans between cortices of primary visual function to cortices involved in complex cognitive functions such as control, abstract constructs, decision making, memory and (social) cognition, but also (pre)motor areas. Moreover, it captures cortical morphological alterations associated with bipolar disorder, as shown by its correlation to cortical thickness alteration patterns. Our results thus indicate that RC G2 captures variation in receptor similarity that is best summarized as separating unimodal from transmodal cortices. The relevance of NTRM co-expression patterns to differentiate sensory from association areas is in line with recent work that employed principal component analysis to autoradiography-derived NTRM densities^72^. This correspondence across methodological approaches is especially important, since PET imaging is of considerably lower resolution and cannot pick up on cortical layering as a pivotal determinant of receptor and transporter expression^10^. Furthermore, RC G2 establishes a link between bipolar disorder cortical thinning patterns and receptor fingerprints, notably to 5-HTT, DAT and NMDA co-expression. These receptors have been implicated in genesis and treatment of bipolar disorder^73–76^. As both RC G1 and RC G2 outline meaningful relationships between receptor expression profiles and disease morphology, receptor similarity could provide novel perspectives in the understanding of the neurobiological basis underlying psychiatric diseases. Investigating receptor co-expression patterns rather than focusing on single molecules could shed light on the enigmatic mechanism of actions of psychotropic drugs, especially when taking into account that most take effect through binding multiple types and classes of receptor molecules^77–79^.

Lastly, the third receptome gradient, RC G3, is anchored between the occipital and temporal cortices. It separates GABAa expression patterns from D2, 5-HT1a, CB1, MU, 5-HT4 and VAChT. Its relationship to other modes of brain organization is best described by its significant correlation to the second gradient of structural connectivity. It also significantly correlates to the first gradient of functional connectivity, and gradients of cytoarchitectural variation. Term-based decoding of brain activation reveals that it separates cortices involved in visual processes from cortices involved in auditory processes. RC G3 separates visual from limbic cortices, which differentiates it from the two aforementioned receptome gradients, where limbic and visual cortices are closely aligned. This separation is also described by SC G2. In general, gradient-based analysis indicates that visual and limbic cortices are relevant drivers of cortical receptor similarity axes, as they are polar at either one (RC G1 and G2) or anchors of a gradient (RC G3). Hierarchical clustering of average NTRM densities separate the both the visual and limbic network from other functional networks, mirroring clustering results obtained via autoradiography^80^, and indicating more unique chemoarchitectural compositions in these regions. Summarizing the interrelationships of receptome gradients and brain structure and function, our results suggest that receptor similarity is organized in a fashion that combines organizational principles of cytoarchitectural, structural and functional differentiation, although interrelationships to structural and functional connectivity and cytoarchitectural variation present differently across parcellation granularities. Incorporating receptor similarity as a novel layer in studies of structure-function relationships might thus be crucial in discerning a governing set of rules in hierarchical brain architecture^81^.

Analysis of architectural correspondence on the node-level showed significant decoupling of SC and FC from RC in particular in heteromodal and paralimbic regions, whereas primary areas showed strongest coupling. This suggests that both structure-function as well as structure-structure relationships dissociate in regions conveying more abstract cognitive processes such as attention, cognitive control, and memory^82–85^. Previous work has shown that structural and functional connectivity are more closely linked in unimodal cortices and exhibit gradual decoupling towards transmodal cortices, a phenomenon that is hypothesized to be instrumental for human flexible cognition^86–88^. Replicating this observation for receptor similarity suggests that diversification of receptor fingerprints may be equally important to enable flexible cognitive functions^1^. We corroborate this hypothesis through clustering analysis, where we found that functional networks involved in higher-order cognitive functions and heteromodal cortices show greater receptomic diversity, meaning a wider spread of receptor fingerprints represented in them. This is consistent with the observation that associative areas show high segregation into sub-areas based on their receptor architecture^89^. High receptomic diversity might be a disease vulnerability factor, as recent work has shown that cortical thickness alterations across different diseases are most pronounced in heteromodal cortices^90^. Notably, our results exemplify a chemoarchitectural divide between heteromodal and paralimbic cortices, as the latter show similar receptor co-distribution homogeneity to idiotypic cortices. A mechanistic explanation might be that, next to memory and emotion^91^, olfactory areas are also located in paralimbic cortices, which adds a sensory component to their function^92^. Additionally, recent work has indicated a differentiation between heteromodal and paralimbic regions, where the former show decreased heritability and cross-species similarity^88^. We thus argue for a more nuanced differentiation of paralimbic from heteromodal cortices, as increasing evidence of architectural differences challenges viewing their relationship purely through a unimodal-to-transmodal lens.

Finally, our analysis of subcortical regions provides novel insights into the chemoarchitecture of subcortical structures and their projections to the cortex. Hierarchical agglomerative clustering of NTRM fingerprints reveals a meaningful separation of subcortical structures based on their functionality, exemplified by the differentiation of striatal structures (putamen, accumbens and caudate nuclei) and pallidal globe from thalamus. Striatum and pallidal globe constitute the basal ganglia, which, together with the thalamus, form the cortico-basal ganglia-thalamic loop. Here, basal ganglia are implicated in motor functions and complex signal integration, while the thalamus orchestrates the communication between large-scale cortical networks^93–95^. This functional divide is not only reflected in receptor fingerprints, but also in receptomic Leiden clustering and principal gradient decomposition, where the first principal subcortical receptomic gradient describes a striato-thalamic axis. We thus expand a chemoarchitecturally-driven structure-function relationship observed in the cortex^15–17,80^ to subcortical structures. Furthermore, we observe partial similarity in receptor co-expression patterns driving subcortical and cortical receptor similarity. While differences in co-expression patterns of 5-HT4 and M1 from α4β2 and NAT seem to be relevant in both cortex and subcortex, the two areas differ in other relevant co-expression patterns. For example, 5-HTT and α4β2 distributions in the cortex are prominently anticorrelated but show similar distributions in the subcortex. Irrespective of individual receptor co-expressions, a general similarity in subcortical and cortical receptome organization is indicated by overlapping cortical and subcortico-cortical receptome gradients. Considering similarities and differences in receptor fingerprints could be important when investigating the modulating influence of subcortico-cortical projections on functional brain networks ^95,96^.

It is of note that the resource we used to comprise the receptome, while extensive, does not contain all cerebral neurotransmitters. Important molecules such as the α2 noradrenaline receptor, which is an important drug target in the central nervous system^97,98^, are missing from our dataset. Our findings must be viewed with the incompleteness of our primary resource in mind. Furthermore, while PET scans were performed on healthy participants, information on medication and medical history was not available for all participants. Thus, we cannot control for potential medication or disease effects. Additionally, the comparatively low spatial resolution of PET imaging is exacerbated by the group-average nature of our dataset. This especially limits the ability to investigate subcortical structures. For example, the thalamus consists of more than 60 nuclei with distinct cellular composition and diverging functionality^99^, important properties we cannot pick up on. Other important subcortical structures, e.g. the subthalamic nuclei, cannot be confidently studied due to their size, limiting our whole-brain perspective to larger nuclei. A more detailed analysis of the subcortical receptome will require methods with higher resolution^100^. Additionally, due to the normalization of tracer uptake in PET images to the cerebellum, our resource also does not permit the analysis of receptor similarity in the cerebellar cortex, limiting our analyses to the telencephalon.

In sum, our work outlines the organization of receptor similarity across the cortex and subcortical structures, yielding an additional layer of brain organization that has meaningful connections to brain structure and function in both health and disease. Incorporating this layer in future studies may provide important steps towards answering the question of how flexible cognition is supported by its physical substrates. Meeting this ultimate goal will provide new avenues to understand, treat and prevent psychiatric diseases and lessen both the personal and societal burden posed by mental illnesses.

## Materials & Methods

### NTRM data generation

To investigate cortical and subcortical NTRM covariance, we made use of an open-access dataset described previously^45^. The associated receptor/transporters, tracers, number of healthy participants, ages, and original publications, for which we refer to full methodological details, are listed in **Table S1**. In brief, images were acquired in healthy participants, using best practice imaging protocols recommended for each radioligand^101^ and averaged across participants before being shared. Images were registered to the MNI152 template (2009c, asymmetric). No medication history of participants was available. The accuracy and validity of receptor density as derived from the PET images has been confirmed using autoradiography data, and mean age of participants was shown to have negligible influence on tracer density values^45^. The cortical receptor density maps were parcellated to 100, 200, 300 and 400 regions based on the Schaefer parcellation^51^, averaging the intensity values per parcel. Subcortical NTRM densities were extracted using a functional connectivity-derived topographic atlas^102^. For tracers where more than one study was included, a weighted average was generated. This resulted in a parcel x 19 matrix of format (parcel x receptor). The intensity values were z-score normalized per tracer. We then performed parcel x parcel Spearman rank correlation of receptor densities, yielding the receptome matrix of regional receptor similarity.

### Principal gradient decomposition

To assess the driving axes of cortical and subcortical covariance organization, we employed principal gradient decomposition^65^ using the brainspace python package^103^. To calculate principal gradients of cortical NTRM covariance, rsFC and MPC, the full matrix was used. SC gradients were separately calculated for intrahemispheric connections in both hemispheres, using procrustes analysis to align the gradients to increase comparability, and subsequently concatenated. To calculate the principal gradients, the respective input matrices were thresholded at 90% and, using a normalized angle similarity kernel, transformed into a square non-negative affinity matrix. We then applied diffusion embedding^104^, a nonlinear dimensionality reduction technique, to extract a low-dimensional embedding of the affinity matrix. Diffusion embedding projects network nodes into a common gradient space, where their distance is a function of connection strengths. This means that nodes closely together in this space display either many supra-threshold or few very strong connections, while nodes distant in gradient space display weak to no connections. In diffusion embedding, a parameter α controls the influence of sampling density on the underlying manifold (where α = 0 equals no influence and α = 1 equals maximal influence). Similar to previous work^65^, we set α to 0.5 to retain global relations in the embedded space and provide robustness to noise in the original matrix.

### Structural, functional and microstructural profile covariance data generation

To contextualize receptor similarity organization, we aimed to compare it to structural connectivity (SC), resting state functional connectivity (FC) and Microstructural Profile Covariance (MPC). The diversity pertaining to age and sociodemographic variables of the subjects in the PET dataset made the selection of matched reference subjects for FC, SC and MPC analysis infeasible. Instead, we opted for the construction of group-consensus FC, SC and MPC matrices collected from the same healthy individuals, obtained and processed in a reproducible pipeline to ultimately provide comparability of the receptome to SC, FC and MPC measures of reference nature. We thus chose the Microstructure Informed Connectomics (MICA-MICs) dataset^105^ to obtain FC, SC and MPC data. MRI data was acquired at the Brain Imaging Centre of the Montreal Neurological Institute and Hospital, using a 3T Siemens Magnetom Prisma-Fit equipped with a 64-channel head coil, from 50 healthy young adults with no prior history of neurological or mental illnesses (23 women; 29.54±5.62 years). No medication history was available. For each participant, (1) a T1-weighted (T1w) structural scan, (2) multi-shell diffusion-weighted imaging (DWI), (3) resting-state functional MRI (rs-fMRI), and (4) a second T1-weighted scan, followed by quantitative T1 (qT1) mapping. Image preprocessing was performed via micapipe, an open-access processing pipeline for multimodal MRI data^106^. Individual functional connectomes were generated by averaging rs-fMRI time series within cortical parcels and cross-correlating all nodal time series. Individual structural connectomes were defined as the weighted count of tractography-derived whole-brain streamlines. To estimate individual microstructural profile covariance, 14 equivolumetric surfaces were generated to sample vertex-wise qT1 intensities across cortical depths, and subsequently averaged within parcels. Parcel-level qT1 intensity values were cross-correlated using partial correlations while controlling for the average cortical intensity profile. The resulting values were log-transformed to obtain the individual MPC matrices^47^.

To generate the group-average matrix of each modality, precomputed and pre-parcellated matrices of 50 individual subjects were used. As no PET data was available for the medial wall, the rows and columns representing it in all SC, FC and MPC matrices were discarded. For SC and FC matrices additionally, rows and columns containing values for subcortical regions were discarded as well, as no analysis of subcortical SC and FC was intended. To generate the group-consensus MPC matrix, parcel values across the subjects were averaged. To generate the group-consensus FC matrix, the subject matrices underwent Fisher’s r-to-z transformation, and subsequently, parcel values across the subjects were averaged. To generate the group-consensus SC matrix, individual matrices were log-transformed and parcel values across subjects were averaged. Afterwards, we applied distance-dependent thresholding to account for the over-representation of short-range and under-representation of long-range connections in non-thresholded group-consensus SC matrices^107^, and the resulting thresholded matrix was used in subsequent analyses.

### Coupling analysis

To investigate the coupling between receptor similarity and FC, SC and MPC, we performed row-wise Spearman rank correlation analyses of the non-zero elements of the respective matrices.

### Leiden clustering

To evaluate whether receptor similarity intrinsically structures the cortical surface and subcortical structures, we applied the Leiden clustering algorithm^52^. The Leiden algorithm is a greedy optimization method that aims to maximize the number of within-group edges and minimize the number of between-group edges, with the resulting network modularity being governed by the resolution parameter ɣ. To incorporate anticorrelations, we used a negative-assymetric approach, meaning that we aimed to maximize positive edge weights within communities and negative edge weights between communities. To search the feature space, we chose a ɣ range of 0.5 to 10 in increments of 0.05 for cortical data, calculating 1000 partition solutions per ɣ. For subcortical structures, we chose a ɣ range of 1 to 10 in increments of 0.5, calculating 250 partitions per ɣ. To assess partition stability, we calculated the z-rand score for every partition with every other partition per ɣ value and chose the partition with the highest mean z-rand score, indicating highest similarity to all other partitions for the given ɣ^108,109^. Additionally, we calculated the variance of z-rand scores between partitions per ɣ. A high mean z-rand score and a low z-rand score variance indicated a stable partition solution.

### Modular stability

To assess the overlap of cortical partitions derived from rsFC and receptomic clustering, we developed the modular stability score. The modular stability score quantifies to what degree functional networks break up into different receptomic communities with an increased receptomic network modularity. It is calculated as 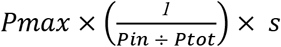, where Pmax is the biggest single partition in a network, Pin is the number of partitions inside the network, Ptot is the total number of partitions, and s is the network size in percent.

### Meta-analytic decoding

To assess the functional loadings of principal receptome gradients, we performed correlation analyses with meta-analysis derived terms of brain activation from the neurosynth database^49^. We parcellated pre-computed activation maps supplied in the brainstat toolbox^110^ and correlated the term-specific activation patterns with principal receptome gradient values, resulting in a total of 3328 values of gradient-term correlations, which were subsequently tested for statistical significance. We then extracted correlation strengths of terms that cover topics from unimodal to transmodal functionality, as well as neurological and psychiatric diseases. A full list of terms can be found in (List supplement). In cases where redundancy was present (e.g. “disorder asd” and “autism spectrum”), the strongest correlating version of the term was selected.

### Disorder impact

To assess the relationship between principal receptome gradients and various neurological and psychiatric diseases, we used publicly available multi-site summary statistics of cortical thinning published by the ENIGMA Consortium^50^. Covariate-adjusted case-vs-control differences, denoted by across-site random-effects meta-analyses of Cohen’s d-values for cortical thickness, were acquired through the ENIGMA toolbox python package^53^. Multiple linear regression analyses were used to fit age, sex, and site information to cortical thickness measures. Before computing summary statistics, raw data was preprocessed, segmented and parcellated according to the Desikan-Killiany atlas in FreeSurfer (http://surfer.nmr.mgh.harvard.edu) at each site and according to standard ENIGMA quality control protocols (see http://enigma.ini.usc.edu/protocols/imaging-protocols). To assess a diverse range of cerebral illnesses, we included eight diseases in our analysis: autism spectrum disorder (ASD)^54^, attention deficit hyperactivity disorder (ADHD)^55^, bipolar disorder (BPD)^56^, DiGeorge-syndrome (22q11.2 deletion syndrome) (DGS)^57^, epilepsy (EPS)^58^, major depressive disorder (MDD)^59^, obsessive compulsive disorder (OCD)^60^ and schizophrenia (SCZ)^61^. Sample sizes ranged from 1,272 (ADHD) to 9,572 (SCZ). Summary statistics were derived from adult samples, except for ASD, where all age ranges were used.

### Hierarchical clustering

To discern a similarity hierarchy of subcortical structures based on mean NTRM density, we performed agglomerative hierarchical clustering. Initially, a set of n samples consists of m clusters, where m=n. In an iterative approach, the samples that are most similar are combined into a cluster, where after each iteration, there are m - # iteration clusters^111^. This process is repeated until m = 1. We use euclidean distance to assess the distance between clusters, and use the WPGMA method to select the closest pair of subsets^112^.

### Null models

Assessment of statistical significance in brain imaging data may be biased when not accounting for spatial autocorrelation of brain imaging signals^113,114^. To generate permuted brain maps that preserve spatial autocorrelation in parcellated data, we resorted to variogram matching (VGM)^115^. Here, we randomly shuffle the input data, and then apply distance-dependent smoothing and rescaling to recover spatial autocorrelation. To assess the significance when comparing surface-projected data, we applied spin permutation^113^ to generate randomly permuted brain maps by random-angle spherical rotation of surface-projected data points, which preserves spatial autocorrelation. Parcel values that got rotated into the medial wall, and values from the medial wall that got rotated to the cortical surface, were discarded^116^. In each approach, we generated 1000 permuted brain maps.

## Data availability

All data and software used in this study is openly accessible. PET data is available at https://github.com/netneurolab/hansen_receptors. FC, SC and MPC data is available at https://portal.conp.ca/dataset?id=projects/mica-mics. ENIGMA data is available through enigmatoolbox (https://github.com/MICA-MNI/ENIGMA). Meta-analytical functional activation data is available through brainstat (https://github.com/MICA-MNI/BrainStat). The code used to perform the analyses can be found at https://github.com/CNG-LAB/cngopen/receptor_similarity.

## Supporting information

Supplemental List 1

## Acknowledgements

We thank Nicola Palomero-Gallagher and Thomas Funck for helpful discussion. JYH acknowledges support from the Helmholtz International BigBrain Analytics & Learning Laboratory and the Natural Sciences and Engineering Research Council of Canada. BM acknowledges support from the Natural Sciences and Engineering Research Council of Canada (NSERC), Canadian Institutes of Health Research (CIHR), Brain Canada Foundation Future Leaders Fund, the Canada Research Chairs Program, the Michael J. Fox Foundation, and the Healthy Brains for Healthy Lives initiative. BCB acknowledges research support from the National Science and Engineering Research Council of Canada (NSERC Discovery-1304413), the CIHR (FDN-154298, PJT-174995), SickKids Foundation (NI17-039), Azrieli Center for Autism Research (ACAR-TACC), BrainCanada (Future Leaders), FRQ-S, and the Tier-2 Canada Research Chairs program. SBE acknowledges support from the Human Brain Project. SLV was supported by Max Planck Gesellschaft (Otto Hahn award). BCB, SBE, and SLV were furthermore funded in part by Helmholtz Association’s Initiative and Networking Fund under the Helmholtz International Lab grant agreement InterLabs-0015, and the Canada First Research Excellence Fund (CFREF Competition 2, 2015-2016) awarded to the Healthy Brains, Healthy Lives initiative at McGill University, through the Helmholtz International BigBrain Analytics and Learning Laboratory (HIBALL). This project has received funding from the European Union’s Horizon 2020 research and innovation program under grant agreement No. 826421, “TheVirtualBrain-Cloud”. Last, we acknowledge and thank those who openly shared their data with the neuroscientific community, which enabled us to perform our study, including the PIs involved in the PET scanning, the MICA-MICS dataset, the Neurosynth tool, as well as all members the ENIGMA consortium working groups.

## Supplementary Materials

**Fig. S1:**
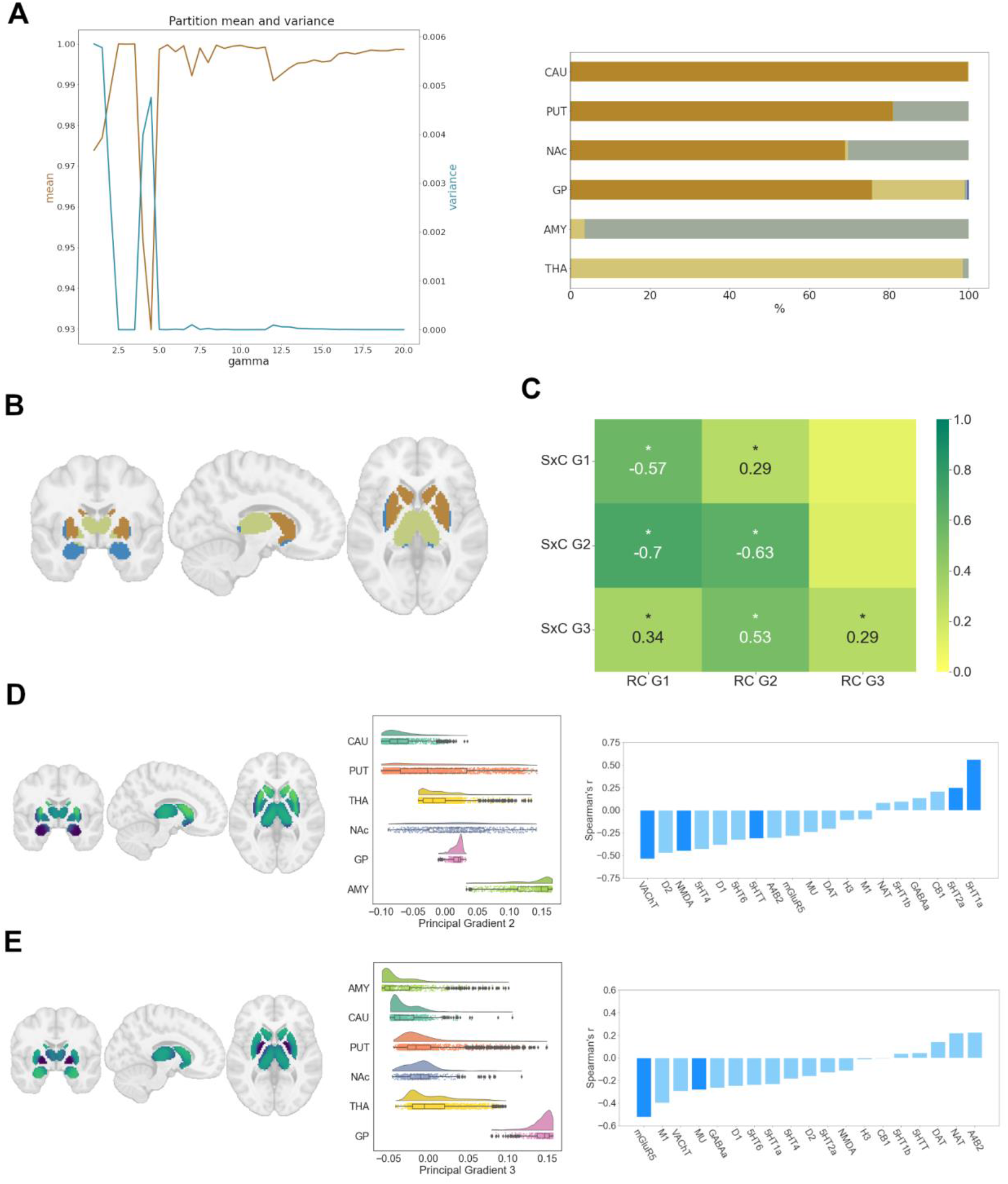
Subcortical receptome. **A)** Leiden clustering of the subcortical receptome. *Left:* Mean and variance of z-rand scores across Leiden algorithm partition resolutions. Note that the clustering results are very stable across gammas. *Right:* Distribution of a stable partition at gamma=2.5 across subcortical structures. **B)** Projection of a stable partition at gamma=2.5 to subcortical structures. **C)** Spearman rank correlation of subcortico-cortical to cortical receptome gradients. **D)** Principal receptome gradient decomposition of the subcortical receptome. *Left:* Subcortical projection of the second principal gradient of the subcortical receptome. *Middle:* Distribution of sRC G2 values across subcortical structures. *Right:* Spearman rank correlations of sRC G2 values with individual NTRM densities. **E)** Principal receptome gradient decomposition of the subcortical receptome. *Left:* Subcortical projection of the third principal gradient of the subcortical receptome. *Middle:* Distribution of sRC G3 values across subcortical structures. *Right:* Spearman rank correlations of sRC G3 values with individual NTRM densities.

**Fig. S2.**
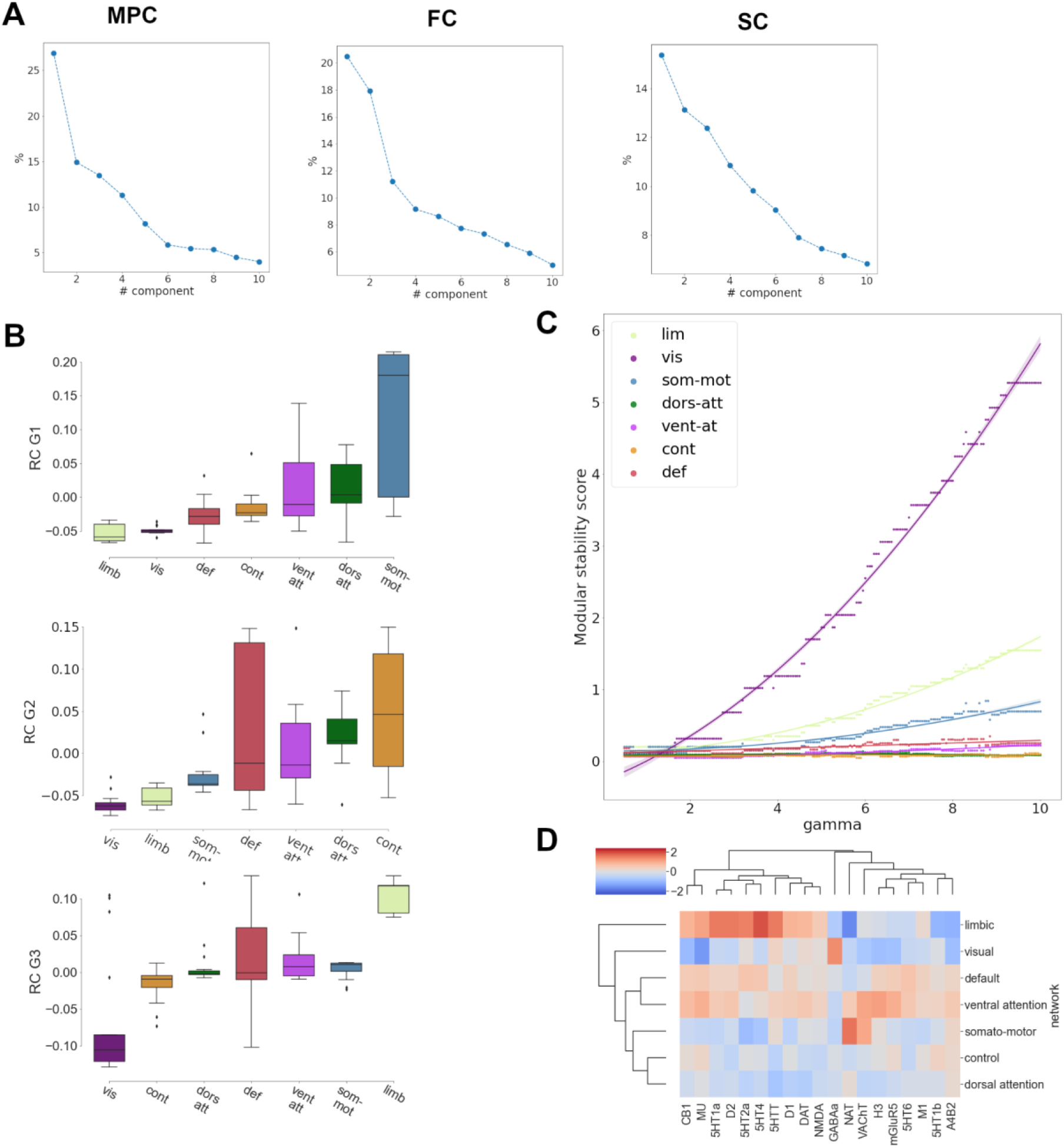
Contextualization of gradients in hierarchical brain organization. **A**) Variance explained by principal gradient decomposition. *Left:* MPC; *Middle:* FC; *Right:* SC **B**) Distribution of RC G1 (*top*), RC G2 (*middle*) and RC G3 (*bottom*) values across functional networks **C**) Modular stability of receptome similarity clustering in functional networks **D**) Hierarchical clustering, using euclidean distance and the WPGMA algorhithm, of NTRM densities in functional networks.

**Fig. S3.**
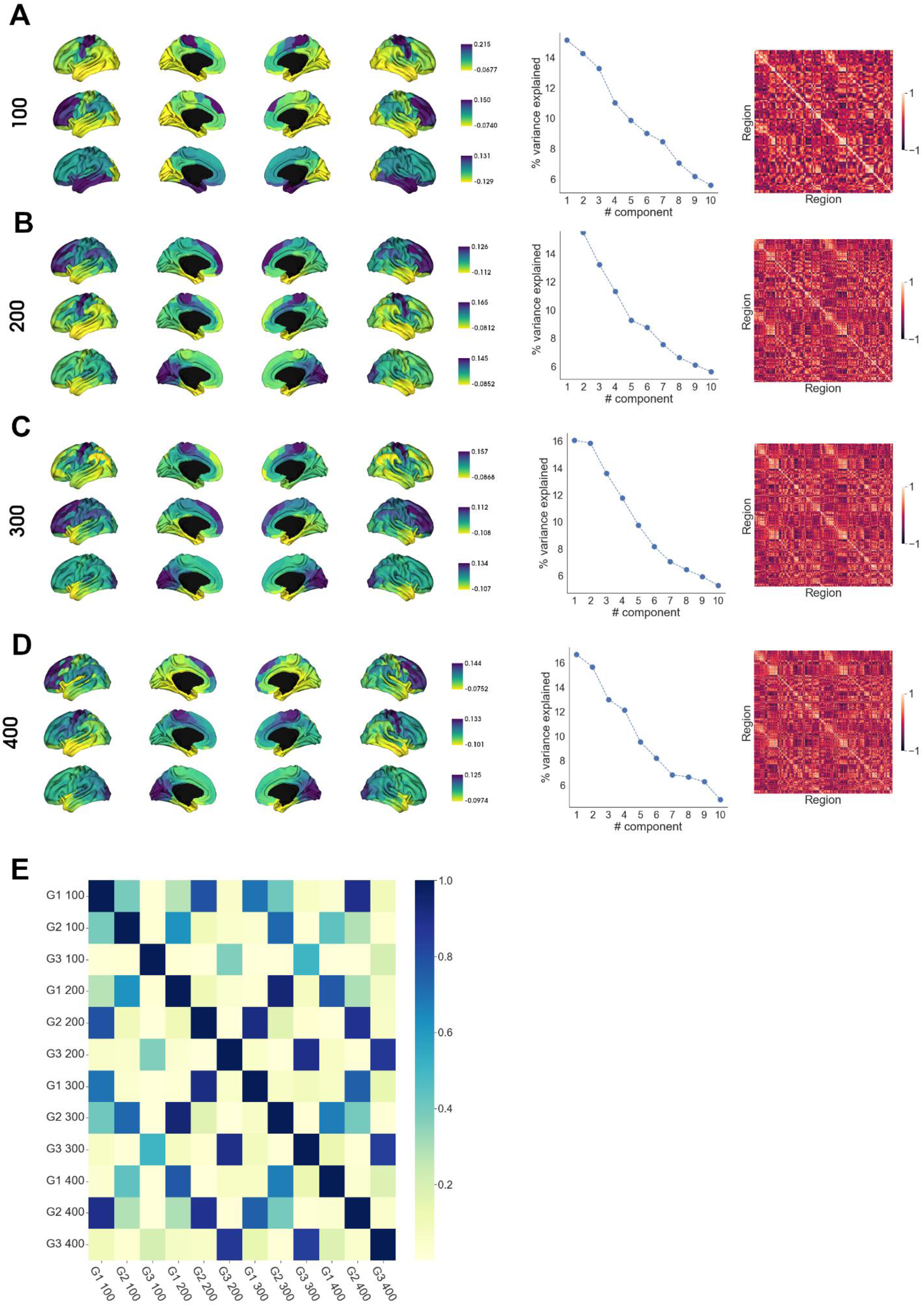
Robustness of receptome gradients and functional decoding. Robustness of principal receptome gradient decomposition across different parcellation granularities. *Left:* RC G1, RC G2 and RC G3 (top-to-bottom) projected on the cortical surface. *Middle*: Variance explained by principal gradient decomposition. *Right:* Receptome matrix. **A**) 100 parcels, **B**) 200 parcels, **C**) 300 parcels, **D**) 400 parcels. **E**) Squared Spearman rank correlation coefficients between neurosynth terms-receptome gradient values associations.

**Fig. S4.**
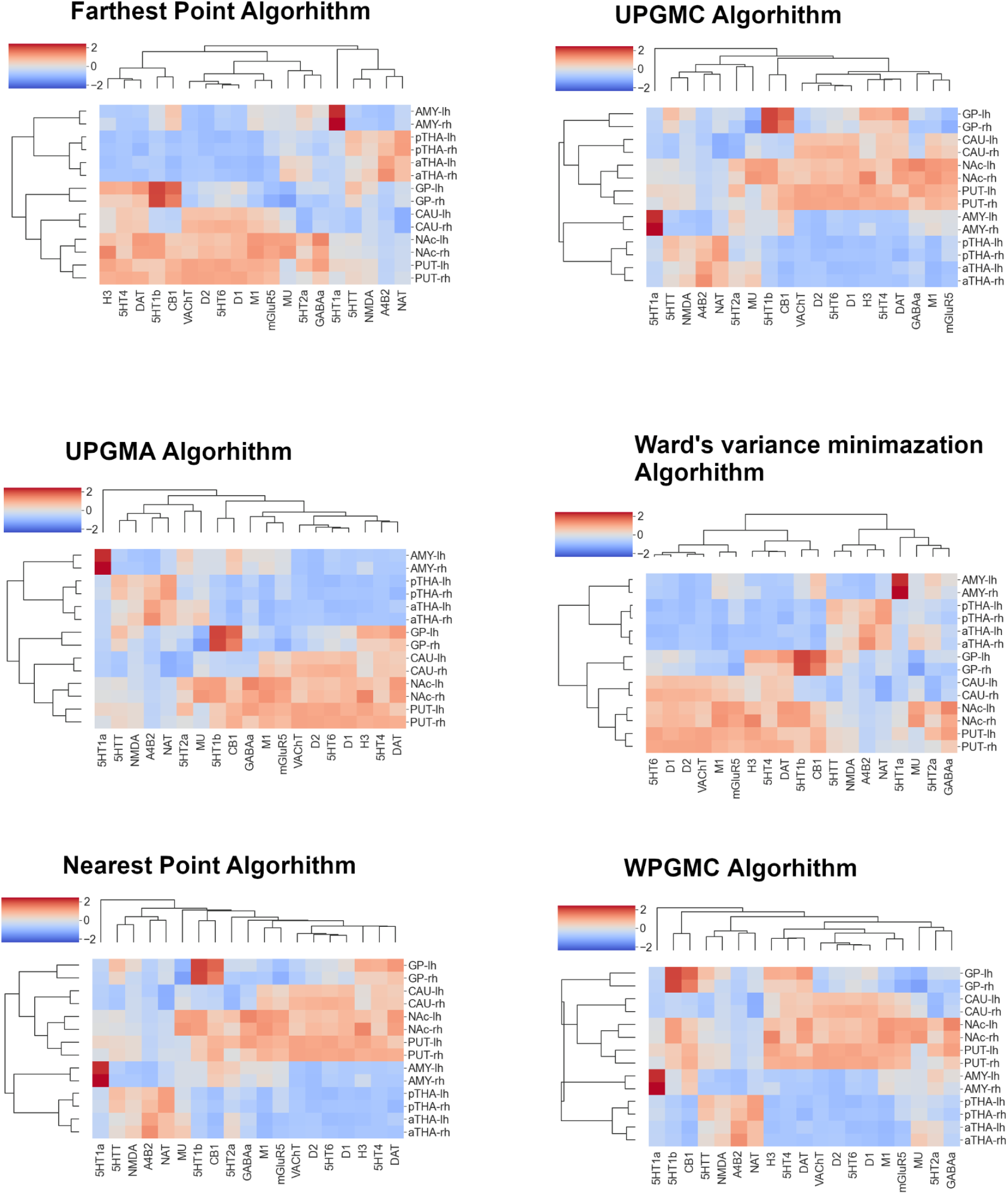
Robustness of agglomerative hierarchical clustering - subcortex. Replication of agglomerative hierarchical clustering of average NTRM densities in subcortical structures, using euclidean distance and different linkage methods.

**Fig. S5.**
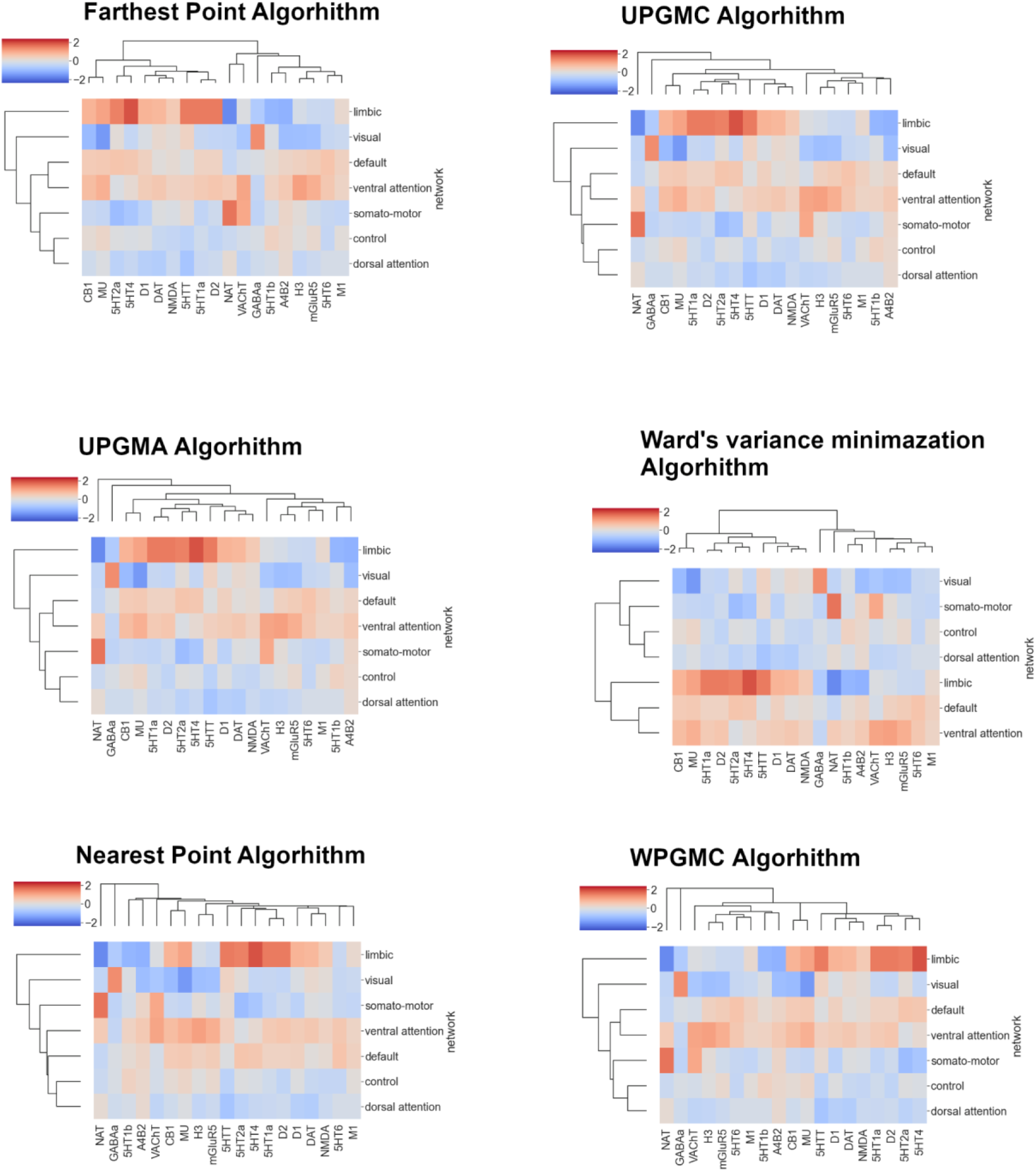
Robustness of agglomerative hierarchical clustering - cortex. Replication of agglomerative hierarchical clustering of NTRM densities in functional networks, using euclidean distance and different linkage methods.

**Table S1:** Neurotransmitter receptors and transporters included in analyses. BPND = non-displaceable binding potential; VT = tracer distribution volume; Bmax = density (pmol/ml) converted from binding potential or distributional volume using autoradiography-derived densities; SUVR = standard uptake value ratio. Neurotransmitter receptor maps without citations refer to unpublished data. Table adapted from Hansen et al^45^

| NTRM | Neurotransmitter | Tracer | Measure | N | Age | References |
| --- | --- | --- | --- | --- | --- | --- |
| **D1** | dopamine | [11C]SCH23390 | BPND | 13 | 33 ± 13 | Kaller et al., 2017<sup>117</sup> |
| **D2** | dopamine | [11C]FLB-457 | BPND | 37 | 48.4 ± 16.9 | Smith et al., 2019<sup>118,119</sup> |
| **D2** | dopamine | [11C]FLB-457 | BPND | 55 | 32.5 ± 9.7 | Sandiego et al., 2015<sup>118–122</sup> |
| **DAT** | dopamine | [123I]-FP-CIT | SUVR | 174 | 61 ± 11 | Dukart et al., 2018<sup>123</sup> |
| **NET** | norepinephrine | [11C]MRB | BPND | 77 | 33.4 ± 9.2 | Ding et al., 2010<sup>124–127</sup> |
| **5-HT1A** | serotonin | [11C]WAY-100635 | BPND | 36 | 26.3 ± 5.2 | Savli et al., 2012<sup>128</sup> |
| **5-HT1B** | serotonin | [11C]P943 | BPND | 65 | 33.7 ± 9.7 | Gallezot et al., 2010<sup>129–135</sup> |
| **5-HT1B** | serotonin | [11C]P943 | BPND | 23 | 28.7 ± 7.0 | Savli et al., 2012<sup>128</sup> |
| **5-HT2A** | serotonin | [11C]Cimbi-36 | Bmax | 29 | 22.6 ± 2.7 | Beliveau et al., 2017<sup>18</sup> |
| **5-HT4** | serotonin | [11C]SB207145 | Bmax | 59 | 25.9 ± 5.3 | Beliveau et al., 2017<sup>18</sup> |
| **5-HT6** | serotonin | [11C]GSK215083 | BPND | 30 | 36.6 ± 9.0 | Radhakrishnan et al., 2018<sup>136,137</sup> |
| **5-HTT** | serotonin | [11C]DASB | Bmax | 100 | 25.1 ± 5.8 | Beliveau et al., 2017<sup>18</sup> |
| **α4β2** | acetylcholine | [18F]flubatine | VT | 30 | 33.5 ± 10.7 | Hillmer et al., 2016<sup>138,139</sup> |
| **M1** | acetylcholine | [11C]LSN3172176 | BPND | 24 | 40.5 ± 11.7 | Naganawa et al., 2021<sup>140</sup> |
| **VACHT** | acetylcholine | [18F]FEOBV | SUVR | 4 | 37 ± 10.2 | PI: Lauri Tuominen & Synthia Guimond |
| **VACHT** | acetylcholine | [18F]FEOBV | SUVR | 18 | 66.8 ± 6.8 | Aghourian et al., 2017<sup>141</sup> |
| **VACHT** | acetylcholine | [18F]FEOBV | SUVR | 5 | 68.3 ± 3.1 | Bedard et al., 2019<sup>142</sup> |
| **NMDA** | glutamate | [18F]GE-179 | VT | 29 | 40.9 ± 12.7 | Galovic et al., 2021<sup>143–145</sup> |
| **mGluR5** | glutamate | [11C]ABP688 | BPND | 73 | 19.9 ± 3.04 | Smart et al., 2019<sup>146</sup> |
| **mGluR5** | glutamate | [11C]ABP688 | BPND | 22 | 67.9 ± 9.6 | PI: Pedro Rosa-Neto & Eliane Kobayashi |
| **mGluR5** | glutamate | [11C]ABP688 | BPND | 28 | 33.1 ± 11.2 | DuBois et al., 2016<sup>147</sup> |
| **GABAA** | GABA | [11C]flumazenil | Bmax | 16 | 26.6 ± 8 | Nørgaard et al., 2021<sup>19</sup> |
| **H3** | histamine | [11C]GSK189254 | VT | 8 | 31.7 ± 9.0 | Gallezot et al., 2017<sup>148</sup> |
| **CB1** | cannabinoids | [11C]OMAR | VT | 77 | 30.0 ± 8.9 | Normandin et al., 2015<sup>149–152</sup> |
| **MU** | opioid | [11C]carfentanil | BPND | 204 | 32.3 ± 10.8 | Kantonen et al., 2020<sup>153</sup> |

